# Clinical proteomics reveals vulnerabilities in non-invasive breast ductal carcinoma and drives personalized treatment strategies

**DOI:** 10.1101/2023.07.11.548580

**Authors:** Georgia Mitsa, Livia Florianova, Josiane Lafleur, Adriana Aguilar-Mahecha, Rene P. Zahedi, Sonia V del Rincon, Mark Basik, Christoph H Borchers, Gerald Batist

**Affiliations:** Division of Experimental Medicine, McGill University, Montreal, QC; Segal Cancer Proteomics Centre, Lady Davis Institute for Medical Research, Jewish General Hospital, Montreal, QC; Department of Pathology, Segal Cancer Centre, Lady Davis Institute for Medical Research, Jewish General Hospital, McGill University, Montreal, QC; Segal Cancer Centre, Lady Davis Institute for Medical Research, Jewish General Hospital, Montreal, QC; Manitoba Centre for Proteomics and Systems Biology, Winnipeg, MB; Department of Internal Medicine, University of Manitoba, Winnipeg, MB; Department of Oncology and Surgery, Lady Davis Institute for Medical Research, Jewish General Hospital, McGill University, Montreal, QC; Department of Oncology, McGill University, Montreal, QC; Exactis Innovation, Montreal, QC

**Author notes:** **Correspondence** Gerald Batist, MD, Lady Davis Institute for Medical Research, Jewish General Hospital, McGill University, Montréal, Quebec, H3T 1E2, Canada, Christoph H Borchers, Ph.D., Segal Cancer Proteomics Centre, Lady Davis Institute for Medical Research, Jewish General Hospital, McGill University, Montréal, Quebec, H3T 1E2, Canada.

## Abstract

Ductal carcinoma in situ (DCIS) is the most common type (80%) of noninvasive breast lesions. The lack of validated prognostic markers, limited patient numbers and variable tissue quality significantly impact diagnosis, risk stratification, patient enrolment, and results of clinical studies. We performed label-free quantitative proteomics on 50 clinical formalin-fixed, paraffin embedded biopsies, validating 22 putative biomarkers from independent genetic studies. Our comprehensive proteomic phenotyping reveals more than 380 differentially expressed proteins and metabolic vulnerabilities, that can inform new therapeutic strategies for DCIS and IDC. Due to the readily druggable nature of proteins and metabolites, this study is of high interest for clinical research and pharmaceutical industry. To further evaluate our findings, and to promote the clinical translation of our study, we developed a highly multiplexed targeted proteomics assay for 90 proteins associated with cancer metabolism, RNA regulation and signature cancer pathways, such as Pi3K/AKT/mTOR and EGFR/RAS/RAF.

## 1 Introduction

Ductal carcinoma in situ (DCIS) is a pre-invasive (stage 0) neoplastic lesion that is associated with a ∼10-fold elevated risk of developing invasive breast cancer, e.g., invasive ductal carcinoma (IDC).^1^ Due to this increased risk, patients diagnosed with DCIS undergo aggressive treatment with breast conserving surgery or total mastectomy with optional adjuvant therapy, i.e., radiation or endocrine therapy.

Studies, however, show that if left untreated, only 20-50% of DCIS patients will progress to IDC.^2-5^ This has led to global concerns regarding overtreatment of DCIS patients, the resulting high economic burden for the healthcare system and, most importantly, a high psychological burden for the patients. Tools and expression signatures to predict invasive progression for better informed clinical decision making are required and many international trials are currently enrolling patients with DCIS for non-surgical management by active surveillance, e.g., LORIS, LORD and LARRIKIN.^6^ The COMET trial (NCT02926911) in the US is targeting histologically confirmed low-risk DCIS for a comparison of surgery to monitoring and endocrine therapy.

At present, the diagnosis of DCIS is based on calcifications observed during mammography screenings and histological assessment of tissue biopsies, i.e., formalin-fixed and paraffin embedded (FFPE) needle core biopsies. Five morphological key features, high intra-tumor heterogeneity, poor inter-observer agreement,^7-10^ and the lack of validated prognostic markers significantly impact clear diagnosis and risk stratification, as well as patient enrolment and final results of clinical studies.

There is currently no precision oncology treatment available for patients diagnosed with DCIS. Post-operative (adjuvant) therapy is guided by immunohistochemistry (IHC) assays for estrogen and progesterone receptor status (ER and PR), HER2 expression status (by fluorescence in situ hybridization, FISH), as well as BRCA1/2 mutation status. Clinical multigene assays, such as Oncotype DX/DCIS, MammaPrint or PreludeDx DCIS, are sometimes used to clinically predict recurrence risks of patients but are not standard and only guide the use of adjuvant therapy.

Generally, DCIS studies are limited by patient number and tissue quality. Recent genomic landscaping studies on individual DCIS lesions identified putative biomarkers associated with progression towards IDC and give insights into the underlying cancer biology. Multi-omic profiling of DCIS, however, is still challenging since DCIS and IDC lesions are mostly studied in FFPE-preserved samples; pure DCIS lesions can be very small in size as they are usually from minimally invasive needle core biopsies, and access to pure IDC lesions is limited, as most surgically removed IDC lesions also present in-situ components and may follow effective neoadjuvant therapy.

The current study makes use of our recently published FFPE-proteomics method that facilitates proteomic profiling on FFPE-preserved tissue cores.^11^ In a cohort of carefully curated patients treated with DCIS and IDC at the Segal Cancer Centre of the Jewish General Hospital (JGH) in Montreal (n=51) we investigate changes in the protein expression of 29 pure DCIS lesions, 18 pure IDC lesions, 13 mixed-type lesions (IDC with in-situ components), and 9 cases where pure DCIS and pure IDC is present in different lesions in the same patient, either synchronously or metachronously (see Fig. 1). Data from recently published independent gene expression studies investigating the progression from DCIS to IDC were used to complement the label-free protein expression data. Since FFPE preservation eliminates up to 85% of metabolites,^12-16^ we used Quantitative Systems Metabolism (QSM™) technology from Doppelganger Biosystem GmbH, Germany, an AI-driven metabolic analysis using proteomics data,^17^ for a comprehensive profiling of the central metabolism/energy metabolism. Guided by these results, we developed a highly multiplexed parallel-reaction monitoring (PRM) assay for precise quantitation of 90 proteins, that are associated with cancer metabolism, RNA regulation and major cancer growth-associated pathways, such as PI3K/AKT/mTOR and EGFR/RAS/RAF.

**Figure 1:**
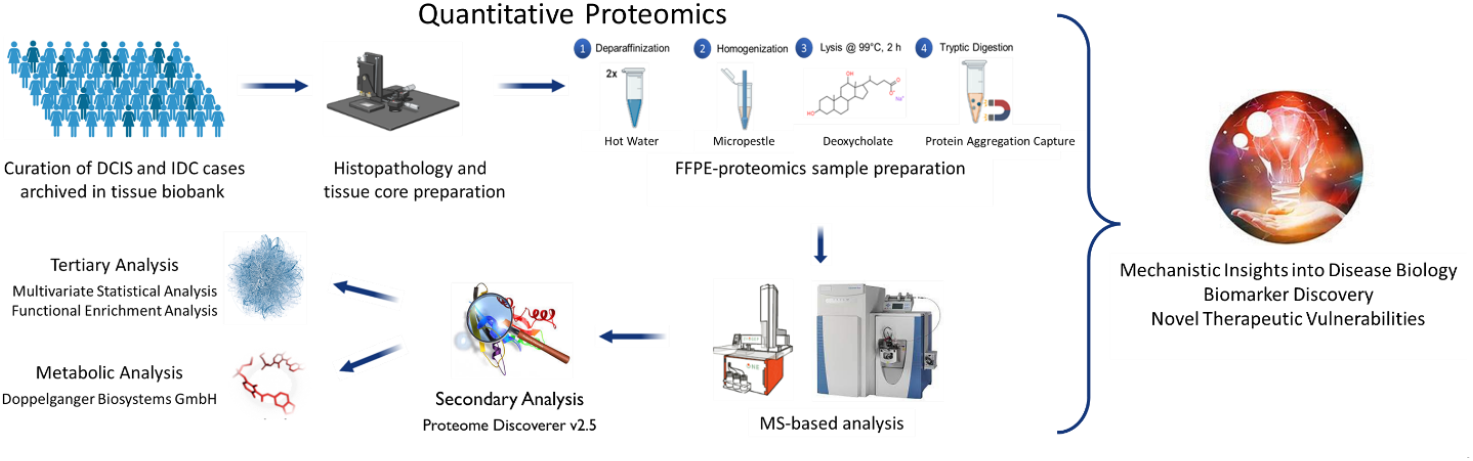
Experimental Design. Label-free quantitative proteomics was performed in a cohort of carefully curated patients treated with DCIS and IDC (n=51) to investigate changes in the protein expression of 29 pure DCIS lesions, 18 pure IDC lesions, 13 mixed-type lesions (IDC with in-situ components), and 9 cases where pure DCIS and pure IDC is present in different lesions in the same patient, either synchronously or metachronously. The protein extraction of FFPE tissue cores (1 mm diameter, ∼0.8 mm^3^ tissue volume) used an optimized FFPE-proteomics protocol published here.^11^ The samples were analyzed on ‘plug-and-play’ platform built for standardization in clinical proteomics, and the data was processed using state-of-the-art data analysis tools, including machine learning/AI-driven algorithms for improved and higher confidence mechanistic insights.

## 2 Results

### 2.1 DCIS and IDC are highly heterogeneous tumor phenotypes but build two distinct clusters in sparse Partial Least Squares Regression for Discrimination Analysis (sPLS-DA)

Several genomic centered studies have reported that both DCIS and IDC tumor phenotypes are highly heterogeneous,^8-10,18,19^ hampering clinical diagnosis but also limiting statistical power and robust assay development to complement clinical diagnosis. Using a streamlined FFPE-proteomics workflow ^11^ with a standard label-free mass spectrometry (MS)-based data analysis, we quantified more than 2800 proteins at a 1% false discovery rate (FDR) on the protein and peptide level. Using less than 1% of the total protein extracted from a single 1-mm FFPE tissue core, we cover 6 orders of magnitude of the DCIS/IDC proteome (Fig. 2a). Notably, the proteome of the two ductal breast cancer disease states seems to be clearly differential from each other, as an sPLS-DA shows two distinct clusters between the study cohorts (Fig. 2b). The sPLS-DA is a statistical method used for extracting and selecting important features from high-dimensional data to discriminate between different groups, while simultaneously considering sparsity to improve interpretability and reduce overfitting.^20^ Based on the available clinical data (non-omics data) and small sample size, we are not in the position to infer any underlying patterns or biological relationships leading to this clustering on the protein level. Nevertheless, the top 10 features driving the proteomic variability between DCIS and IDC seem to reflect high transcriptional activity, extracellular matrix remodeling and inflammation processes (Figs. 2c and 2d).

**Figure 2:**
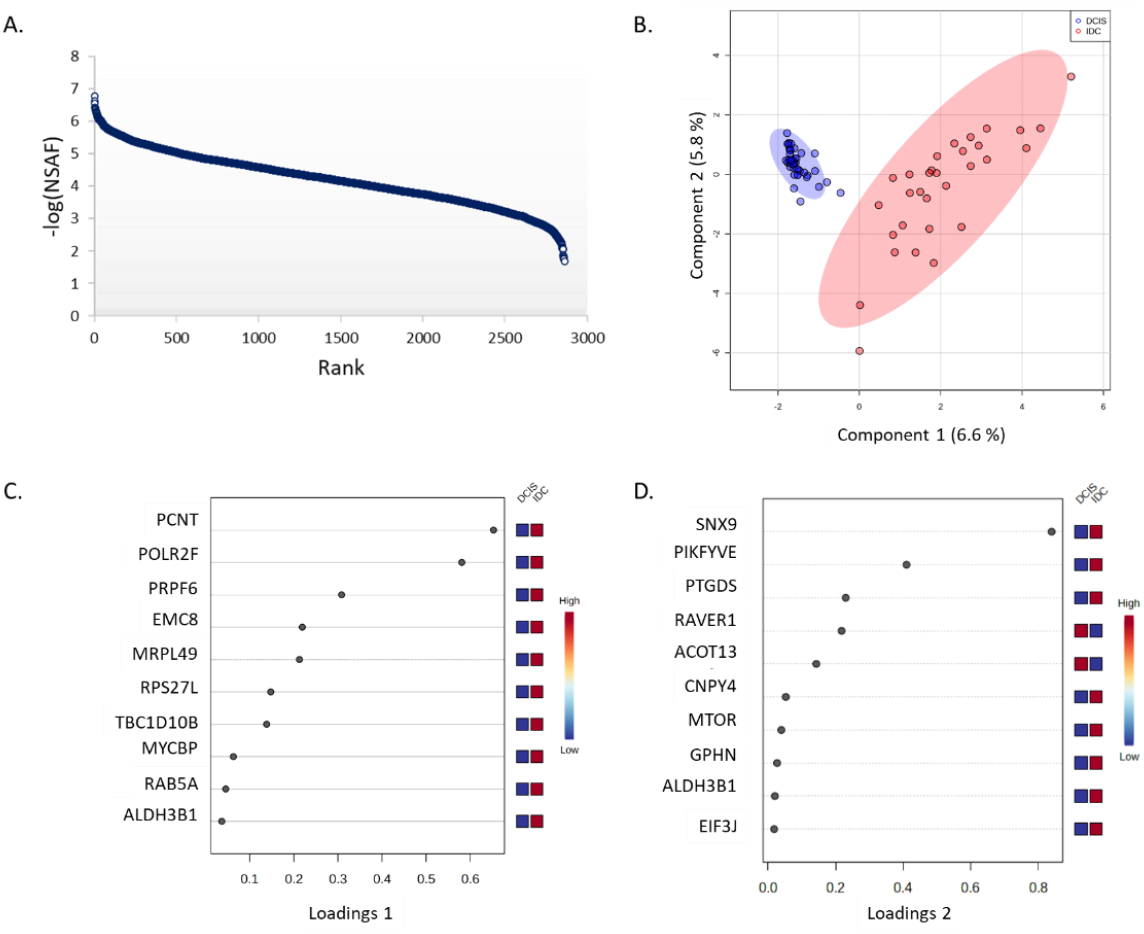
Data Quality and Evaluation of Variability. **(A)** Dynamic range of ∼2,860 proteins quantified in ductal breast cancer, at a 1% false discovery rate. All –log10 values were based on normalized spectral abundance factor (NSAF) values, which were used to normalize the spectral count. High NSAF values represent a high level of expression. 6 orders of magnitude of the DCIS/IDC proteome are covered using ∼1% of the total sample and a standard data dependent acquisition (DDA) method without fractionation. **(B)** Sparse Partial Least Squares Regression for Discrimination Analysis (sPLS-DA) showing good clustering of the two study groups. The oval shape represents 95% confidence intervals. Interquartile and ROUT method identified no outlier samples. **(C/D)** Loading plots of the sPLS-DA, showing proteins/genes that drive the variability and clustering between DCIS and IDC. The right x-axis shows expression levels of these drivers in the DCIS/IDC samples.

### 2.2 MS-based proteomics complements and supports independent genomic/transcriptomic studies of DCIS to IDC progression

Study of progression of DCIS to IDC has mainly used gene expression analysis or IHC/FISH on the protein level. Recent studies have demonstrated significant misalignment between genome and even transcriptome and the ultimate protein levels, and IHC is poorly quantitative.^21-24^ Therefore, we sought to confirm these findings using direct measurement of proteins. We compared MS-based label-free proteomics data with 49 differentially expressed genes identified by three recent larger-scale independent genomics/transcriptomics studies ^9,25,26^ and found 22 overlapping genes (see Table 1). Proteomics data identified gene products of *FOXA1, POSTN, THBS2, CA12, FN1* and *ALDH1* as differentially expressed proteins (DEPs, unpaired t-test, p<0.05).

**Table 1:**
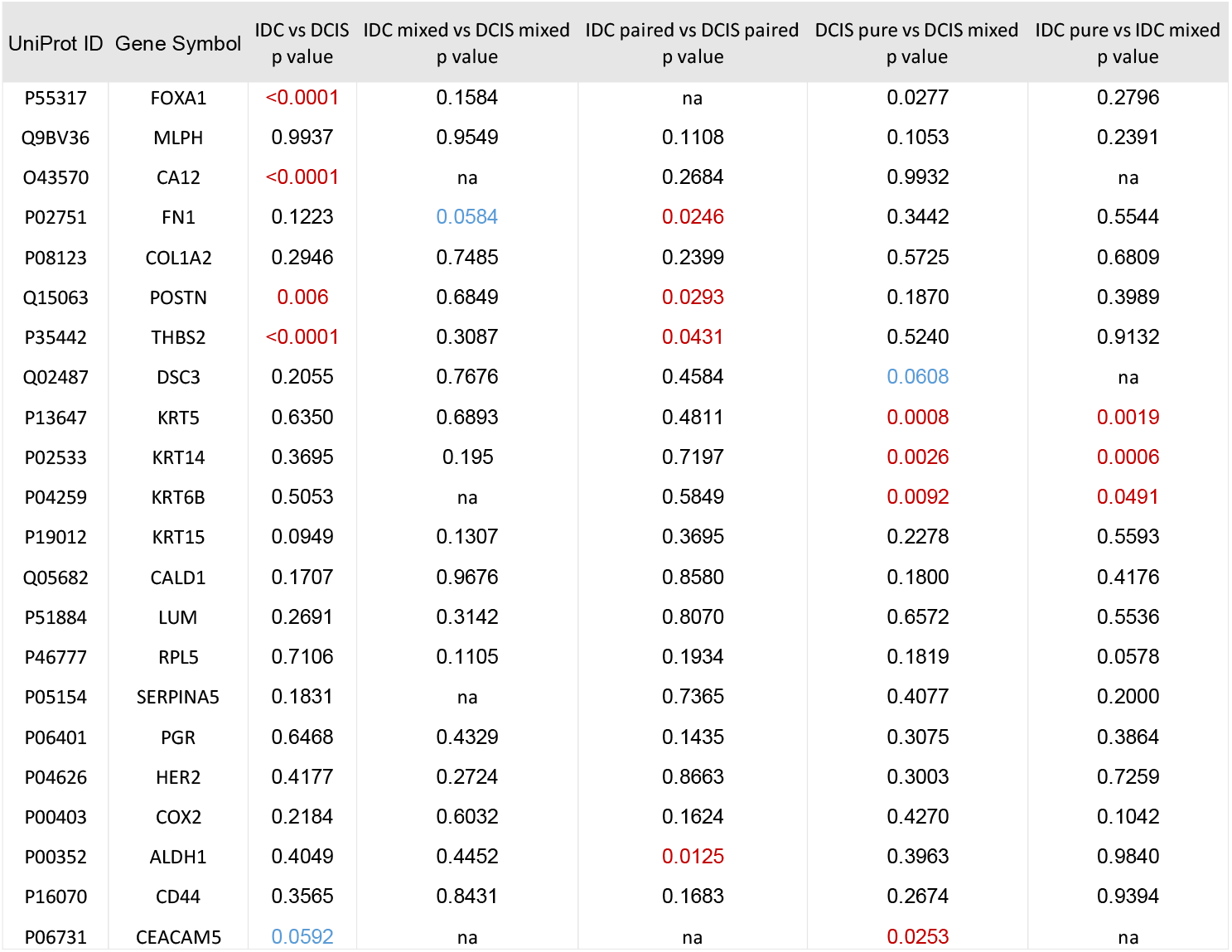
Overlapping molecules from independent gene expression studies and this proteomic profiling. Differential expression values of 22 proteins corresponding to genes proposed in the literature as biomarkers for DCIS to IDC progression. In red statistically significant entities with a student t-test p-value <0.05, in yellow entities close to the set p-value. na = ‘not applicable’; the protein was not quantified in that dataset.

The proteomics data shows lower *FOXA1* (Forkhead Box A1) expression in pure DCIS compared to pure IDC (p<0.0001), and increased expression in mixed-type DCIS compared to pure DCIS (p=0.03) suggesting a protective function of FOXA1. The loss or silencing of *FOXA1* observed in DCIS seems to promote cell migration and invasion. Interestingly, forced expression of *FOXA1* in MCF-7 (IDC cell line) inhibits growth, and controls cell plasticity by repressing the basal-like phenotype.^27,28^ Genetic studies associate *FOXA1* with heterochromatin remodeling, particularly affecting hormone receptor transcription,^29^ and regulation of the cell cycle with *BRCA1*.^30,31^ Evidence of FOXA1 involvement in tumor progression on the (epi-)genetic, transcriptomic, and proteomic level warrants further investigation of FOXA1 as clinical biomarker and its clinical utility for DCIS risk stratification.

POSTN (Periostin), THBS2 (Thrombospondin 2), and FN1 (Fibronectin) mediate cell-cell and cell-matrix interactions. POSTN, a downstream effector of β-catenin, activates PI3K/AKT and ERK pathways.^32^ In DCIS, these proteins have lower expression levels compared to IDC (p<0.03, p<0.04, p<0.03, respectively), indicating stromal remodeling in DCIS to IDC progression.

*CA12* (Carbonic Anhydrase 12) regulates the tumor microenvironment and metabolic pathways,^33-35^ with lower protein levels in pure DCIS compared to pure IDC (p<0.0001). Loss of CA12 activity likely creates a more favorable environment for malignant cell growth and progression towards IDC.

High *ALDH1* (Aldehyde Dehydrogenase 1) expression characterizes cancer stem cells associated with tumorigenesis, metastatic behavior, and poor outcomes.^36,37^ While an IHC-based profiling of DCIS did not associate ALDH1 with breast cancer events,^9^ our MS-based analysis on paired DCIS/IDC lesions, does show a significantly higher concentration of ALDH1 in DCIS compared to IDC lesions (p=0.01), supporting findings from stem cell biology that ALDH1 might be a functional and prognostic biomarker of tumorigenesis in DCIS.

Having access to ‘real-world’ mixed-type lesions, the most prevalent clinical phenotype of breast ductal carcinoma, we were in the unique position to investigate the proteome of DCIS lesions that are likely active in the transition to IDC, depleted from inter-tumor heterogeneity. Comparing pure DCIS to mixed-type DCIS lesions revealed significantly lower protein levels of KRT5, KRT14, KRT6B, and CEACAM5 in pure DCIS lesions (p<0.05), indicating stromal remodeling as a key feature in the progression from pre-cancer to invasive cancer, with prognostic value for DCIS management. High expression of KRTs is linked to good prognosis in breast cancer, while lower levels are associated with invasive tumor proliferation.^38-40^ CEACAM5 (also CEA) expression has context-dependent impact and a protective function in breast cancer, with potential usefulness in disease monitoring.^9,41,42^ Similarly, comparing pure IDC to mixed-type IDC lesions showed a loss of KRT expression in mixed-type IDC (p<0.05), suggesting a protective role of KRTs and marker of progressiveness in DCIS.

### 2.3 Loss of basal membrane stability, inflammatory processes, and epithelial-to-mesenchymal transition (EMT) identified as key events driving DCIS progression

Having confirmed the results of genomic/transcriptomic studies in this setting using direct MS-based protein measurements, we turned to a global proteomics approach to discover further features of the DCIS-IDC scenario.

Differential expression analysis of more than 2800 proteins identified in pure DCIS compared to IDC, revealed ∼388 DEPs using an unpaired t-test with post-hoc Benjamini-Hochberg FDR method for multiple hypothesis testing (q<0.01) and at least a 2-fold-change in protein expression between DCIS and IDC (Fig. 3 a, and Supplemental Table 1). To reduce the inter-patient variability, we compared proteomic profiles of DCIS and IDC lesions from the same patients (n=9). Ten differentially expressed proteins (DEPs) were identified: ILK, ITGA4, GPRC5A, FNTA, SCPEP1, EPB41L3, and SORBS1 were upregulated in DCIS, while ACAP1, ATP6V0A1, and KPRP were upregulated in IDC (Fig 3 b, and Supplemental Table 2).

**Figure 3:**
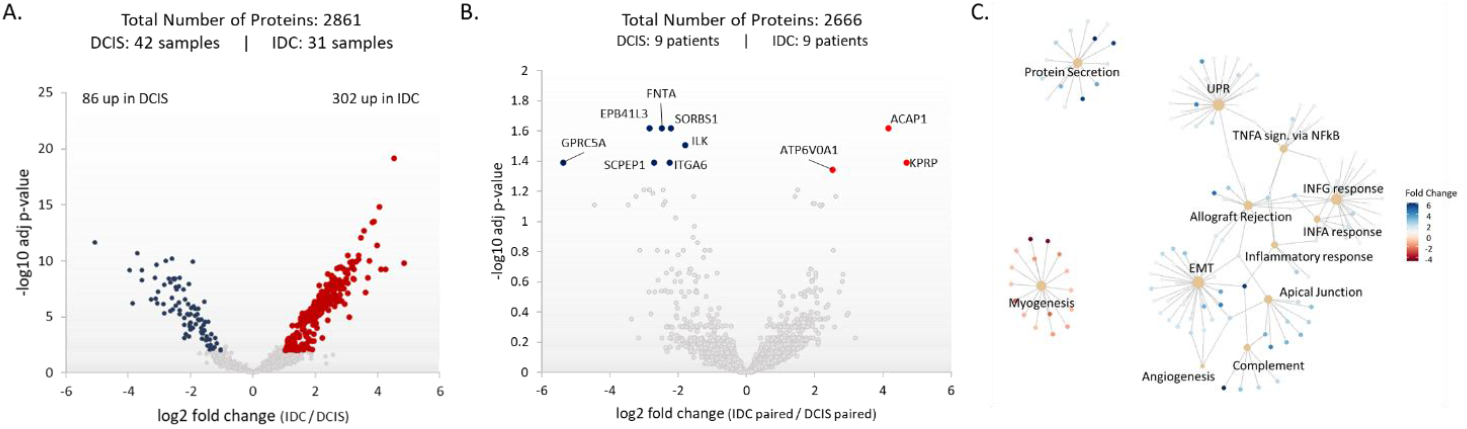
Differential Expression Analysis reflects loss of basal membrane stability, inflammatory processes, and epithelial-mesenchymal transition as key events towards DCIS to IDC progression. (A) Volcano plot of the proteome of pure IDC compared to pure DCIS lesions showing 388 differentially expressed proteins (unpaired t-test with post-hoc Benjamini-Krieger analysis p<0.01, abs log2 fold change >2). (B) Volcano plot of the proteome of paired IDC lesions compared to paired DCIS lesions showing 10 differentially expressed proteins (unpaired t-test with post-hoc Benjamini-Krieger analysis p<0.05, abs log2 fold change >2). (C) Molecular networks representing up-/downregulated pathways in IDC compared to DCIS lesions. UPR: Unfolded Protein Response, EMT: Epithelial-to-Mesenchymal Transition.

**Figure 4:**
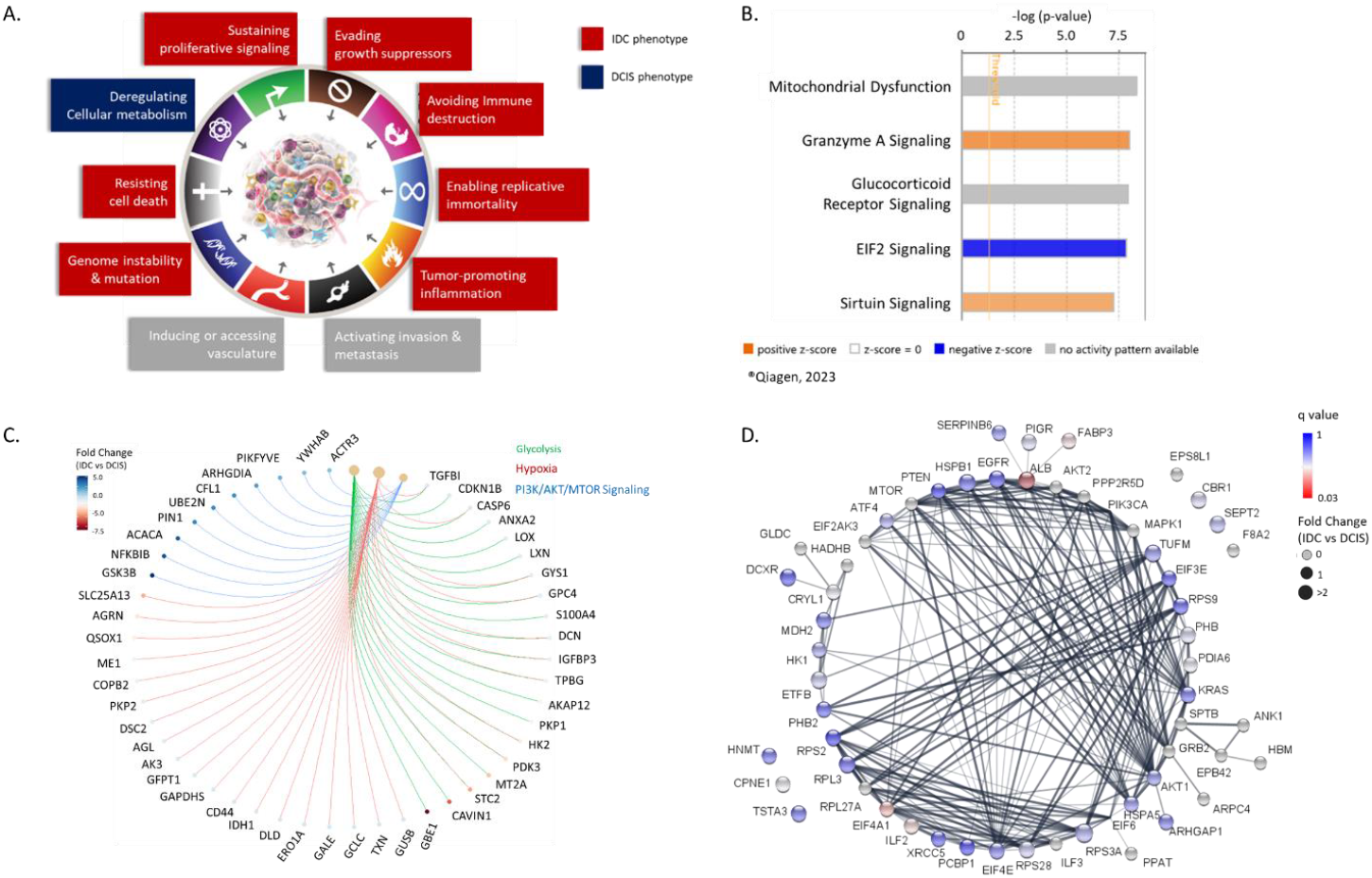
Dysregulation of central energy metabolism is a key event in the DCIS tumor phenotype. (A) Graphical representation of hallmarks of cancer (modified from ^81^) characteristic for proteomic tumor profiling of DCIS and IDC tumors. (B) Top 5 canonical pathways from Ingenuity Pathway Analysis on differentially expressed proteins in 42 DCIS and 31 IDC tumors. (C) Signature proteins potentially driving DCIS progression through glycolysis, hypoxia (or “pseudo-hypoxia”) and PI3K/AKT/mTOR pathway, identified by gene set enrichment analysis. (D) STRING Network showing the protein expression profile of signature proteins, associated with cancer metabolism, RNA regulation and major cancer pathways, such as PI3K/AKT/mTOR and EGFR/RAS/RAF. Absolute concentration of the proteins was determined by parallel reaction monitoring. The color of the nodes represents q values from multiple-hypothesis testing using unpaired t-tests with post hoc correction using the Benjamini-Krieger FDR method (1% FDR). The node size represents the fold-change. Grey nodes were not quantified, either because no SIS/NAT was available or because there were more than 60% missing values. Edges represent physical and/or functional interaction partners based on the STRING database.

ILK, an integrin linked kinase, regulates integrin signaling and is associated with tumor growth and metastasis.^43,44^ ITGA4 mediates cell-cell adhesions and is linked to cancer progression, inflammatory reactions, and ECM stemness.^45-47^ GPRC5A acts as an oncogene or tumor suppressor in different cancers.^48-50^ Androgen receptor-regulated FNTA enhances KRAS signaling and might be involved in tumorigenesis.^51-56^ SCPEP1 is associated with cancer development, growth and metastasis.^57-59^ EPB41L3 is a tumor suppressor involved in apoptosis and cell cycle regulation.^60-63^ Decreased expression in DCIS was observed for ATP6V0A1, which plays a role in pH homeostasis and tumor cell invasion.^64-66^ ACAP1, which is associated with cell proliferation, migration, and immune infiltration in tumors.^67-69^ Loss of ACAP1 could indicate impaired immune response in IDC progression. KPRP, involved in keratinocyte differentiation,^70,71^ might contribute to invasiveness when its expression is lost in DCIS.

Overall, proteomic profiling of DCIS identified more than 380 putative biomarkers (protein level) to clinically profile DCIS lesions for risk stratification and disease management. The association of the differentially expressed proteins quantified in this study with hallmarks of cancer, such as remodeling of the tumor microenvironment (e.g., ILK, ITGA4, SCPEP1), escape of apoptosis (e.g., ILK, GPRC5A, FNTA, EPB41L3), deregulation of apical junction and energy metabolism (e.g., ATP6V0A1, KPRP, ITGA4), as well as inflammation and immune response processes (e.g., ACAP1, ITGA4) (Fig. 3c), warrants further investigation. Further, most of the identified DEPs are readily druggable and re-purposing of FDA-approved anti-inflammatory drugs and antibiotics pose interesting treatment options for DCIS.

### 2.4 EIF2 and PI3K/Akt/mTOR signaling pathway potentially drive IDC phenotype development through dysregulation of central energy metabolism in cancer

A deeper look into the molecular relationships of all the DEPs we’ve identified by functional enrichment analysis and gene set enrichment analysis, confirms the previously reported loss of basal layer integrity and epithelial to mesenchymal transitions (EMT) as key events supporting IDC. Figure 5a highlights cancer hallmarks that are predominant for the IDC- and DCIS phenotype, highlighting the dysregulation of cell metabolism as a key event in the DCIS-phenotype. Proteomic profiling using MS-based techniques revealed metabolic vulnerabilities in DCIS that can provide insights into tumorigenic metabolic mechanisms, that were missed by genomic/transcriptomic analysis alone.

Functional Enrichment Analysis using IPA identifies mitochondrial dysfunction, granzyme A signaling, glucocorticoid receptor signaling and sirtuin signaling as significantly enriched (p-value of overlap <0.01) in our proteomics dataset, suggesting a dysregulation of glucose metabolism, through a shift from oxidative phosphorylation (i.e., tricarboxylic acid (TCA) cycle) to aerobic glycolysis (Figs. 5b and 5c).^72^ Aerobic glycolysis is also known as Warburg Effect and is characterized by high glucose uptake and glycolytic conversion of glucose to lactate to meet the high energy demands of proliferating cells.^73^ During glycolysis, glucose is converted to pyruvate. Cytosolic pyruvate can either enter the TCA cycle for oxidative phosphorylation (OXPHOS) and ATP-production or be converted to lactate. Under normoxia, the metabolic fate of cytosolic pyruvate, and thus glucose metabolism, is regulated by pyruvate dehydrogenase complex (PDH) and lactate dehydrogenase (LDH), where the PDH reaction is favored.^73,74^ PI3K/AKT signaling can modulate the metabolic fate of pyruvate as an upstream regulator of PDH and LDH, creating “pseudo-hypoxic” conditions that favor pyruvate conversion to lactate. The pivotal role of PI3K/AKT as an upstream regulator in metabolic reprogramming is comprehensively reviewed by Hoxhaj et al.^75^ and involves the interaction with other proliferating signaling pathways, such as MAPK and mTOR. Our proteomic analysis of DCIS identified several differentially expressed molecules involved in glycolysis, hypoxia-mediated reactions and PI3K/AKT/mTOR signaling (Fig. 5c) which warrant further investigation. Metabolomic profiling of FFPE specimens is challenging, because ∼85% of metabolites are washed out during the preservation procedure. To nevertheless gain insights into metabolic changes occurring towards IDC progression, we conducted an AI-based metabolic profiling using QSM™ technology, which is supported by more than 500 publications.^17^ Clear metabolic differences between DCIS/IDC lesions from the same patient (paired DCIS/IDC) were identified, but due to the large variability and small sample size (n=9) metabolic differences between the groups were hard to assess. A multitude of functional markers with direct causal relation to ATP production capacity and utilization of glucose were nevertheless identified (Tbl. 2). These findings confirm the dysregulation of energy metabolism towards IDC progression and suggest that the energy demand of transforming pre-invasive cells (DCIS-phenotype) is mainly achieved by fatty acid metabolism and lactate production.

**Table 1:**
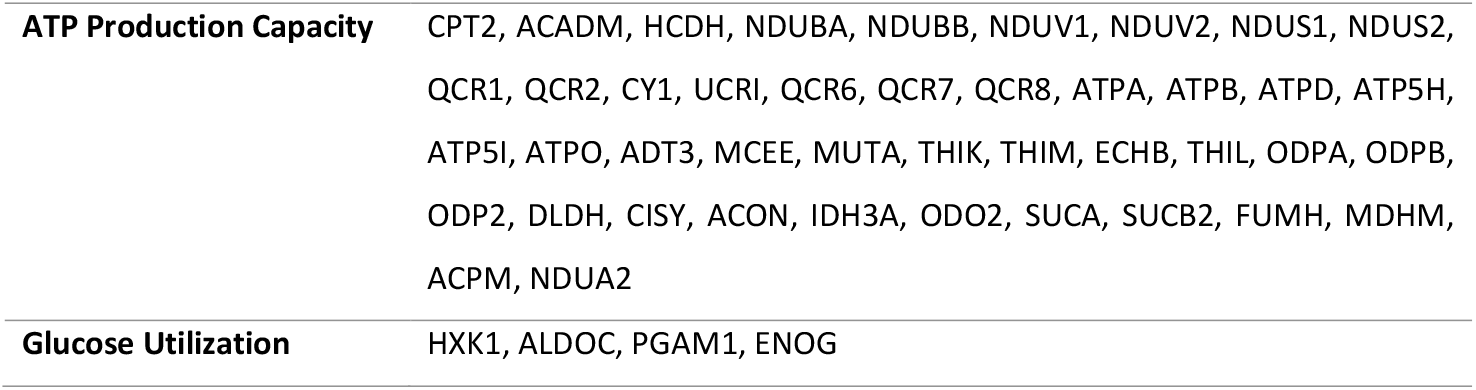
List of putative metabolic biomarkers identified by AI-based metabolic profiling of DCIS and IDC specimens from the same patient.

To further evaluate and promote the translation of our findings into the clinic, we developed a highly multiplexed targeted MS-assay for absolute quantitation of 90 signature peptides, associated with cancer metabolism, central energy metabolism, RNA regulation and members of the PI3K/AKT/mTOR, EIF2 and EGFR/RAS/RAF signaling pathways. A complete list of peptides included in this assay is provided in Supplemental Table 4. The results of the PRM assay are depicted as STRING network (Figure 5d), where the differential expression is represented by the node color, and the absolute fold-change by the node size. These findings correlate well with the previously discussed observations from label-free proteomics and independent genomics/transcriptomics study, showing that DCIS tumors have a tendency towards loss of metabolic functions. Albumin (ALB) is significantly higher expressed in the DCIS phenotype compared to the IDC phenotype (q value = 0.03). Studies associated low albumin levels with changes in the tumor microenvironment to more favorable conditions for disease progression and tumor migration, suggesting that serum albumin levels might have a prognostic value for cancer.^76,77^ Other studies discuss albumin as a potent marker for inflammation and the nutritional status of patients, where low albumin levels correlate with inflammatory processes resulting in higher morbidity and poor prognosis.^78,79^ Our results support these findings, and highlight remodeling of the tumor microenvironment, environmental stress (i.e., malnutrition, which inhibits EIF2 signaling)^80^ and inflammatory processes as key events towards IDC progression.

## 3 Discussion

Clinical research on DCIS has been limited due to low sample numbers, high inter-tumor heterogeneity and low tissue quality, as most DCIS lesions derive from diagnostic needle-core-biopsies and are FFPE embedded. Although genetic/transcriptomic studies of DCIS progression provide a cellular blueprint of what might happen, genes cannot be readily targeted for therapy and post-translational modification cannot be assessed by genetic screening alone. Quantitative proteomics can complement and confirm genetic changes and provides a deeper look into the ‘real-life’ tumor phenotype. The readily druggable nature of proteins makes quantitative proteomics studies attractive for clinical research. Additionally, mass spectrometry-based studies allow both (i) discovery studies for comprehensive tumor profiling and (ii) validation studies in a highly multiplexed manner, with unprecedented accuracy, specificity, and sensitivity.

We established a label-free quantitative proteomics pipeline suitable for needle-core biopsy sized FFPE specimens and performed a comprehensive proteomic phenotyping of DCIS and IDC using less than 1% of the total extracted protein material. We cover 6 orders of magnitude of the disease proteome and identify more than 380 differentially expressed proteins that identify classical hallmarks of cancer, reflective for high transcriptional activity, extracellular matrix remodeling and inflammation processes as key events towards IDC progression. We further identify dysregulation of glucose metabolism as a key event in the transition from pre-invasive to invasive carcinoma. Guided by these results, we developed a highly multiplexed parallel-reaction monitoring (PRM) assay for precise quantitation of 90 proteins, that are associated with cancer metabolism, RNA regulation and major cancer pathways, such as PI3K/AKT/mTOR and EGFR/RAS/RAF. We applied this assay to generate an activation profile of these signature proteins for proliferation and metabolic remodeling in cancer in ‘real world’ clinical samples and were able to support observations from label-free proteomics data with absolute concentrations in the amol range, facilitating the translation of our findings into the clinic. Notably, proteomics profiling revealed that FDA-approved drugs, such as antibiotics and NSAID, may be repurposed for DCIS and IDC treatment, as they have been shown to control and target proteins identified as key events towards IDC progression.

It is important to highlight, that this study design is applicable to many diseases with limited sample volumes and low tissue quality, as it requires only a fraction of the total sample amount allowing discovery and validation studies in the same sample cohort. In our opinion, clinical proteomics is a versatile tool for comprehensive tumor phenotyping, able to capture a ‘real-life’ snapshot of tumor phenotypes, representative of post-translational modifications and epigenetic changes. More than 99% of published clinical biomarkers/genomic assays fail to enter clinical practice,^82^ but we show here that complementing genomics and transcriptomics studies with proteomics data, and vice versa, will help create a better understanding of underlying disease mechanisms and will better inform the selection of biomarker candidates and patient enrolment for clinical studies, ultimately improving the quality and final results of clinical trials.

## 4 Methods

All chemicals and reagents were purchased by Sigma Aldrich (St. Louis, MI, USA) unless otherwise specified. Sequencing grade trypsin (Promega, P/N V511A) was used for the generation of tryptic peptides.

### 4.1 Clinical Specimens

Clinical specimens were obtained from patients who consented for tissue biobanking part of the Jewish General Hospital Breast Biobank (protocol 05-006). The study was performed in accordance with the ethical standards laid down in the 1964 Declaration of Helsinki and was approved by the Jewish General Hospital Research Ethics Board.

A total of 50 clinical cases of patients diagnosed and treated with DCIS and/or IDC at the JGH were carefully curated by a pathologist with expertise in breast cancer to select lesions meeting inclusion criteria for mass spectrometry-based (MS-based) analysis, i.e., at least 30% tumor cellularity and less than 10% necrosis. The patients were of Caucasian ethnicity ranging from 22 to 82 years of age at first diagnosis (median age 52 years). The patients were followed for a period of 1 to 18 years (median 8 years). During the period of follow-up, 43 patients have no evidence of disease, and 1 patient has metastatic disease, while 8 patients died from cancer. The cohort comprises 29 cases with pure DCIS lesions, 18 cases with pure IDC lesions, 13 cases with mixed-type lesions (IDC with in-situ components) and 9 cases with synchronous/metachronous DCIS and IDC. Clinical data for the patients is available upon request.

### 4.2 Sample Preparation

1 mm-diameter tissue cores (∼0.8 mm^3^ tissue volume) were prepared from FFPE-blocks enriching for DCIS or IDC only tumor cells. Excessive paraffin was trimmed off using a clean scalpel blade. Protein extraction was performed following our developed FFPE-proteomics workflow for core needle biopsies. Briefly, paraffin was removed by incubation with hot water (∼80 °C). Each deparaffinized core was mechanically disrupted using a micropestle (Sigma Aldrich, #BAF199230001) in 250 μL of 2% sodium deoxycholate (SDC), 50 mM Tris-HCl, 10 mM tris(2-carboxyethyl)phosphine (TCEP), pH 8.5, followed by sequential incubation on an Eppendorf ThermoMixer C for 20 min at 99 °C (1100 rpm) and for 2 h at 80°C (1100 rpm). Samples were cooled down on ice for 1 minute before a 15-minute centrifugation at 21,000x g (4 °C) to remove cell debris. The supernatant was collected into a Protein LoBinding tube (Eppendorf, Germany) and the total protein concentration was determined using a Pierce Reducing Agent Compatible BCA kit (RAC-BCA, Thermo Scientific, P/N 23252) following the manufacturer’s instructions. Free cysteine residues were alkylated with iodoacetamide to a final concentration of 30 mM and incubation for 30 minutes at room temperature, protected from light.

For 2 μg of protein lysate, 2 μL of ferromagnetic beads with MagReSyn® Hydroxyl functional groups (ReSyn Biosciences, Gauteng, South Africa, 20 μg/mL) were equilibrated with 100 μL of 70% ACN, briefly vortexed and placed on a magnetic rack to remove the supernatant. This step was repeated another two times. Next, the protein extracts were added to the beads and the sample was adjusted to a final concentration of 70% ACN, thoroughly vortexed and incubated for 10 min at room temperature without shaking. The following washing steps were performed on a magnetic rack without disturbing the protein/bead aggregate. The supernatants were discarded, and the beads were washed on the magnetic rack with 1 mL of 95% ACN for 10 s, followed by a wash with 1 mL of 70% ACN without disturbing the protein/bead aggregate. The tubes were removed from the magnetic rack, 100 μL of digestion buffer (1:20 (w/w) trypsin:protein in 0.2 M GuHCl, 50 mM AmBic, 2 mM CaCl_2_) were added and the samples were incubated at 37 °C for 12 h. After acidification with trifluoroacetic acid (TFA) to a final concentration of 2%, the tubes were placed on the magnetic rack for 1 min, followed by removal of the supernatant. To remove residual beads, the samples were centrifuged at 20,000x g for 10 min.

### 4.3 Preparation of spiking solutions for the response curve and absolute quantitation

In order to promote translation of our findings and to validate LFQ Abundances with a more precise targeted MS approach we developed a multiplexed parallel reaction monitoring (PRM) method to quantify 90 proteins in FFPE specimens, measuring the concentration of a unique sig-nature peptide for each protein. All 90 peptides were measured in a single LC-MS/MS run. Two equimolar synthetic peptide mixtures (100 fmol/μg of each peptide) were prepared in 30% ACN with 0.1% formic acid (FA) in water (w/v); one mixture contained unlabeled peptides (light or NAT peptides), and the second mixture contained stable isotope labeled standard peptides (heavy or SIS peptides). The light peptide mixture was used to develop the highly multiplexed PRM assay with optimized peptide-specific parameters, such as collision energy and charge state, while the heavy peptide mixture was used for normalization, serving as spiking solution and internal standard for clinical samples.

Quantitation was performed using a 7-point response curve consisting of a variable amount of light peptides, ranging from 0.41 to 250 fmol (three orders of magnitude), and a constant amount of SIS peptides (50 fmol). Digested bovine serum albumin (BSA, 0.01 μg) was used as surrogate matrix of the response curve. To determine the limit of detection (LOD), a double blank sample was prepared. The blank sample consisted of 0.01 μg BSA digest spiked with 50 fmol of the SIS-mixture and analyzed before and/or directly after the highest calibrant level of the response curve. For quantitation of endogenous protein in the patient samples, 50 fmol of SIS peptide were spiked into 1 μg total digested tissue protein, as determined by RAC-BCA.

### 4.4 Data analysis

1 μg digested protein was pre-concentrated on EV2001 C18 Evotips and separated on a heated (40 °C) EV1137 column (15 cm x 150 μm, 1.5 μm particle size) using Evosep’s “Extended meth-od” (15 samples-per-day (SPD)). The samples were analyzed by data dependent acquisition (DDA) mode, on a Q Exactive Plus Orbitrap mass spectrometer operated with a Nanospray Flex ion source (both from Thermo Fisher Scientific), connected to an Evosep One HPLC (Evosep Bio-systems, Odense, Denmark). Full MS scans were acquired over the mass range from m/z 350 to m/z 1500 at a resolution of 70,000 with an automatic gain control (AGC) target value of 1×106 and a maximum injection time of 50 ms. The 15 most-intense precursor ions (charge states +2, +3, +4) were isolated with a window of 1.2 Da and fragmented using a normalized collision energy of 28; the dynamic exclusion was set to 30 s. MS/MS spectra were acquired at a mass resolution of 17,500, using an AGC target value of 2×104 and a maximum injection time of 64 ms.

Chromatographic separation of all PRM runs was performed with the same equipment and buffers as described above. The Q Exactive Plus was operated in PRM mode at a resolution of 35,000. Target precursor ions were isolated with the quadrupole isolation window set to m/z 1.2. An AGC target of 3×106 was used, allowing for a maximum injection time of 110 ms. Data was acquired in time-scheduled mode, allowing a 2-min retention-time window for each target. Full MS scans were acquired in parallel at low resolution (m/z 17,500) with an AGC target value of 1×106 and a maximum injection time of 50 ms, to ensure sample quality.

MS data files are publicly available through the ProteomeXchange Consortium via the PRIDE partner repository^83^ with the dataset identifier PXD040782. The synthetic peptides selected for this PRM assay were validated by others, information is available through National Cancer Institute’s Clinical Proteomic Tumor Analysis Consortium (CPTAC) Assay Portal (assays.cancer.gov).

### 4.5 Data Processing and Differential Expression Analysis

MS raw data were processed using Proteome Discoverer 2.5 (Thermo Fisher Scientific). Database searches were performed using SequestHT with Multi-Peptide Search (MPS) and a human Swissprot database (January 2019; 20,414 target entries). Label-free quantitation (LFQ) was performed using the Minora feature-detector node within Proteome Discoverer, and the Percolator software was used to calculate posterior error probabilities. Database searches were performed using trypsin as enzyme with a maximum of 2 missed cleavages. Carbamidomethylation of cysteine (+57.021 Da) was set as a fixed modification, and oxidation of methionine (+15.995 Da) as variable modifications. Mass tolerances were set to 10 ppm for precursor ions and 0.02 Da for product ions. The data were filtered to a false discovery rate (FDR) <1% at the peptide and protein levels. Only proteins that were (i) identified with at least one protein unique peptide and (ii) quantified in ≥60% of replicates of at least one of the study groups, were considered for the quantitative comparison. Protein LFQ data obtained from Proteome Discoverer was normalized based on summed protein intensities to correct for differences in sample loading. Missing protein intensity values were imputed using 1.5x the minimum observed intensity for this particular sample. The obtained normalized abundances were used for unpaired t-tests (two tailed, 95% confidence) and differential expression analysis on log2-transformed data with multiple hypothesis testing using the Benjamini-Krieger false-discovery approach (FDR 1%). Proteins having q-values of <0.01 and absolute log2 fold-changes >1 were considered as differential between test-ed groups. Statistical analysis was performed using GraphPad Prism 9 (San Diego, CA, USA).

Raw PRM data were analyzed using Skyline (v22.2.0.351).^84^ Correct peak integration and visual verification of detected peaks was performed manually for each target, and the three to four highest and most stable transitions were selected for quantitation. A linear regression model with 1/x^2^ weighting using the SIS/NAT ratio of each target peptide was used for the calculation of concentrations. Only calibration levels meeting following criteria were accepted for response curve generation and regression analysis; precision average <20% CV per calibration level, accuracy average between 80% and 120% per calibrant level, quantified in at least 3 consecutive calibrant levels. The LOD describes the smallest concentration of the target peptide (analyte) that is likely to be reliably distinguished from instrument noise and at which detection is feasible. To determine the LOD we use replicate injections from a double blank sample, i.e., fixed concentration of the SIS-peptide in surrogate matrix. The average concentration of the double blank plus 3.3x the standard deviation of the blank replicates is used to calculate the lowest detectable concentration for each peptide. The limit of quantitation (LOQ) describes the lowest concentration at which the analyte can not only be reliably detected, but at which above mentioned precision and accuracy criteria are met. Here the LOQ was defined as the lowest calibration level for each peptide. Proteins/Peptides with more than 60% missing values were excluded from downstream analysis.

### 4.6 Functional Enrichment Analysis

Functional Enrichment Analysis was performed using the ‘Core Analysis’ function within Ingenuity Pathway Analysis (Qiagen, Inc., content version: 81348237, release Date: 2022-09-15).^85^ Ingenuity Knowledge Base was used as reference set, allowing direct and indirect relationships. Only molecules having expression p-values <0.05 and absolute log2 fold-changes of >1 were considered for the core analysis. All other settings were kept with default parameters.

### 4.7 Gene Set Enrichment Analysis

A pre-ranked Gene Set Enrichment Analysis (GSEA) was performed using GSEA v4.3.2 (Broad Institute, Inc.) software. The gene list was ranked by differential expression using SIGN function within Excel with calculated log2 fold-change and p-value from an unpaired t-test. A hallmark gene set Molecular Signature Database (MSigDB v2022.1)^86^ was used as references gene set. The search allowed 1000 permutations, with set sizes between 15 and 500 genes. Pathways were collapsed to remove redundancy and to increase selectivity and specificity. Data was visualized using the clusterProfiler^87^ package within R.

### 4.8 Metabolic Profiling

Protein expression data from paired DCIS/IDC cases was sent to Doppelganger Biosystems Inc (Berlin, Germany) for metabolic profiling using Quantitative Systems Metabolism (QSM™) technology.^17^

## Supplemental Materials

The following supporting information can be downloaded at:…. Supplemental Tables T1-T4 with detailed quantitative proteomics data and results from differential expression analysis.

## Author Contributions

Conceptualization, G.M., G.B., C.H.B; methodology, G.M., R.P.Z.; sample preparation and formal analysis, G.M.; writing-original draft preparation, G.M.; writing-review and editing, G.M., A.A.-M., M.B., G.B., C.H.B.; providing of resources, L.F., J.L., A.A-M, M.B., C.H.B. All authors have read and agreed to the published version of the manuscript.

## Acknowledgement

CHB is grateful to Genome Canada for financial support through the Genomics Technology Platform (264PRO). CHB is also grateful for support from the Segal McGill Chair in Molecular Oncology at McGill University (Montreal, Quebec, Canada) and for support from the Terry Fox Research Institute, the Warren Y. Soper Charitable Trust, and the Alvin Segal Family Foundation to the Jewish General Hospital (Montreal, Quebec, Canada). GM would like to acknowledge the patients and their families for their consent to archive biological specimens for clinical research. GM would like to thank Dr. Naciba Benlimame, Lilian Canetti and Dr. James Saliba for their guidance and constructive discussions. This work was done under the auspices of a Memorandum of Understanding between McGill and the U.S. National Cancer Institute’s International Cancer Proteogenome Consortium (ICPC). ICPC encourages international cooperation among institutions and nations in proteogenomic cancer research in which proteogenomic datasets are made available to the public. This work was also done in collaboration with the U.S. National Cancer Institute’s Clinical Proteomic Tumor Analysis Consortium (CPTAC).

## Institutional Review Board Statement

The study was performed in accordance with the ethical standards laid down in the 1964 Declaration of Helsinki and was approved by the Jewish General Hospital Research Ethics Board.

## Informed Consent Statement

Clinical specimens were obtained from patients who consented for tissue biobanking part of the Jewish General Hospital Breast Biobank (protocol 05-006).

## Data Availability Statement

MS data files are publicly available through the ProteomeXchange Consortium via the PRIDE partner repository^83^ with the dataset identifier PXD040782. To access the deposited data visit https://www.ebi.ac.uk/pride/login. Please use *<u>reviewer_pxd040782@ebi</u>*.*<u>ac</u>*.*<u>uk</u>* as the username. The password is *tr3rBy0E*. Clinical data and H&E staining of clinical specimens used in this study are available upon request.

## References

1. Mannu, G.S., et al. Invasive breast cancer and breast cancer mortality after ductal carcinoma in situ in women attending for breast screening in England, 1988-2014: population based observational cohort study. BMJ 369, m1570 (2020).

2. Kumar, A.S., Bhatia, V. & Henderson, I.C. Overdiagnosis and overtreatment of breast cancer: Rates of ductal carcinoma in situ: a US perspective. Breast Cancer Res. 7, 271 (2005).

3. Sanders, M.E., Schuyler, P.A., Dupont, W.D. & Page, D.L. The natural history of low-grade ductal carcinoma in situ of the breast in women treated by biopsy only revealed over 30 years of longterm follow-up. Cancer 103, 2481–2484 (2005).

4. Jones, J.L. Overdiagnosis and overtreatment of breast cancer: Progression of ductal carcinoma in situ: the pathological perspective. Breast Cancer Res. 8, 204 (2006).

5. Collins, L.C., et al. Outcome of patients with ductal carcinoma in situ untreated after diagnostic biopsy: results from the Nurses’ Health Study. Cancer 103, 1778–1784 (2005).

6. Morrissey, R.L., Thompson, A.M. & Lozano, G. Is loss of p53 a driver of ductal carcinoma in situ progression? Br. J. Cancer 127, 1744–1754 (2022).

7. Badve, S., et al. Prediction of local recurrence of ductal carcinoma in situ of the breast using five histological classifications: a comparative study with long follow-up. Hum. Pathol. 29, 915–923 (1998).

8. Badve, S. & Gökmen-Polar, Y. Tumor Heterogeneity in Breast Cancer. Adv. Anat. Pathol. 22, 294–302 (2015).

9. Badve, S.S., et al. Multi-protein spatial signatures in ductal carcinoma in situ (DCIS) of breast. Br. J. Cancer 124, 1150–1159 (2021).

10. Gerdes, M.J., et al. Single-cell heterogeneity in ductal carcinoma in situ of breast. Mod. Pathol. 31, 406–417 (2018).

11. Mitsa, G., et al. A Non-Hazardous Deparaffinization Protocol Enables Quantitative Proteomics of Core Needle Biopsy-Sized Formalin-Fixed and Paraffin-Embedded (FFPE) Tissue Specimens. Int. J. Mol. Sci. 23, 4443 (2022).

12. Buszewska-Forajta, M., et al. Paraffin-embedded tissue as a novel matrix in metabolomics study: optimization of metabolite extraction method. Chromatographia 82, 1501–1513 (2019).

13. Cacciatore, S. & Loda, M. Innovation in metabolomics to improve personalized healthcare. Annals of the New York Academy of Sciences 1346, 57–62 (2015).

14. Cacciatore, S., et al. Metabolic Profiling in Formalin-Fixed and Paraffin-Embedded Prostate Cancer TissuesMetabolic Profile in FFPE Tissues. Mol. Cancer Res. 15, 439–447 (2017).

15. Dannhorn, A., et al. Evaluation of Formalin-Fixed and FFPE Tissues for Spatially Resolved Metabolomics and Drug Distribution Studies. Pharmaceuticals 15, 1307 (2022).

16. Neef, S.K., et al. Optimized protocol for metabolomic and lipidomic profiling in formalin-fixed paraffin-embedded kidney tissue by LC-MS. Anal. Chim. Acta 1134, 125–135 (2020).

17. Berndt, N., Kann, O. & Holzhütter, H.G. Physiology-based kinetic modeling of neuronal energy metabolism unravels the molecular basis of NAD(P)H fluorescence transients. J. Cereb. Blood Flow Metab. 35, 1494–1506 (2015).

18. Nachmanson, D., et al. The breast pre-cancer atlas illustrates the molecular and microenvironmental diversity of ductal carcinoma in situ. NPJ Breast Cancer 8, 6 (2022).

19. Nagasawa, S., et al. Genomic profiling reveals heterogeneous populations of ductal carcinoma in situ of the breast. Commun Biol 4, 438 (2021).

20. Sorochan Armstrong, M.D., de la Mata, A.P. & Harynuk, J.J. Review of Variable Selection Methods for Discriminant-Type Problems in Chemometrics. Frontiers in Analytical Science 2(2022).

21. Jain, A.P., et al. Pan-cancer quantitation of epithelial-mesenchymal transition dynamics using parallel reaction monitoring-based targeted proteomics approach. J. Transl. Med. 20, 1–13 (2022).

22. Chakraborty, S., Hosen, M.I., Ahmed, M. & Shekhar, H.U. Onco-multi-OMICS approach: a new frontier in cancer research. BioMed research international 2018(2018).

23. Dunn, J., et al. Integration and comparison of transcriptomic and proteomic data for meningioma. Cancers (Basel) 12, 3270 (2020).

24. Harnik, Y., et al. Spatial discordances between mRNAs and proteins in the intestinal epithelium. Nature metabolism 3, 1680–1693 (2021).

25. Rebbeck, C.A., et al. Gene expression signatures of individual ductal carcinoma in situ lesions identify processes and biomarkers associated with progression towards invasive ductal carcinoma. Nat Commun 13, 3399 (2022).

26. Dettogni, R.S., et al. Potential biomarkers of ductal carcinoma in situ progression. BMC Cancer 20, 119 (2020).

27. Bernardo, G.M., et al. FOXA1 represses the molecular phenotype of basal breast cancer cells. Oncogene 32, 554–563 (2013).

28. Bernardo, G.M. & Keri, R.A. FOXA1: a transcription factor with parallel functions in development and cancer. Biosci. Rep. 32, 113–130 (2012).

29. Seachrist, D.D., Anstine, L.J. & Keri, R.A. FOXA1: A Pioneer of Nuclear Receptor Action in Breast Cancer. Cancers (Basel) 13(2021).

30. Williamson, E.A., et al. BRCA1 and FOXA1 proteins coregulate the expression of the cell cycle-dependent kinase inhibitor p27(Kip1). Oncogene 25, 1391–1399 (2006).

31. Wolf, I., et al. FOXA1: Growth inhibitor and a favorable prognostic factor in human breast cancer. Int. J. Cancer 120, 1013–1022 (2007).

32. Liu, T., Zhou, L., Xiao, Y., Andl, T. & Zhang, Y. BRAF Inhibitors Reprogram Cancer-Associated Fibroblasts to Drive Matrix Remodeling and Therapeutic Escape in Melanoma. Cancer Res. 82, 419–432 (2022).

33. Barnett, D.H., et al. Estrogen receptor regulation of carbonic anhydrase XII through a distal enhancer in breast cancer. Cancer Res. 68, 3505–3515 (2008).

34. Li, Y., et al. High expression of carbonic anhydrase 12 (CA12) is associated with good prognosis in breast cancer. Neoplasma 66, 420–426 (2019).

35. Ning, W.R., et al. Carbonic anhydrase XII mediates the survival and prometastatic functions of macrophages in human hepatocellular carcinoma. J. Clin. Invest. 132(2022).

36. Charafe-Jauffret, E., et al. Aldehyde dehydrogenase 1-positive cancer stem cells mediate metastasis and poor clinical outcome in inflammatory breast cancer. Clin Cancer Res 16, 45–55 (2010).

37. Douville, J., Beaulieu, R. & Balicki, D. ALDH1 as a functional marker of cancer stem and progenitor cells. Stem Cells Dev. 18, 17–25 (2009).

38. Han, W., Hu, C., Fan, Z.J. & Shen, G.L. Transcript levels of keratin 1/5/6/14/15/16/17 as potential prognostic indicators in melanoma patients. Sci. Rep. 11, 1023 (2021).

39. Saha, S.K., et al. KRT19 directly interacts with beta-catenin/RAC1 complex to regulate NUMB-dependent NOTCH signaling pathway and breast cancer properties. Oncogene 36, 332–349 (2017).

40. Saha, S.K., Yin, Y., Chae, H.S. & Cho, S.-G. Opposing Regulation of Cancer Properties via KRT19-Mediated Differential Modulation of Wnt/β-Catenin/Notch Signaling in Breast and Colon Cancers. in Cancers (Basel), Vol. 11 (2019).

41. Bechmann, M.B., Brydholm, A.V., Codony, V.L., Kim, J. & Villadsen, R. Heterogeneity of CEACAM5 in breast cancer. Oncotarget 11, 3886–3899 (2020).

42. Yang, C., et al. Down-regulation of CEACAM1 in breast cancer. Acta Biochim Biophys Sin (Shanghai) 47, 788–794 (2015).

43. Hannigan, G.E., et al. Regulation of cell adhesion and anchorage-dependent growth by a new beta 1-integrin-linked protein kinase. Nature 379, 91–96 (1996).

44. Delcommenne, M., et al. Phosphoinositide-3-OH kinase-dependent regulation of glycogen synthase kinase 3 and protein kinase B/AKT by the integrin-linked kinase. Proc. Natl. Acad. Sci. U. S. A. 95, 11211–11216 (1998).

45. Kinashi, T. Overview of integrin signaling in the immune system. Methods Mol. Biol. 757, 261–278 (2012).

46. Mo, J., et al. The early predictive effect of low expression of the ITGA4 in colorectal cancer. J. Gastrointest. Oncol. 13, 265–278 (2022).

47. Pulkka, O.P., et al. Clinical relevance of integrin alpha 4 in gastrointestinal stromal tumours. J Cell Mol Med 22, 2220–2230 (2018).

48. Zhou, H. & Rigoutsos, I. The emerging roles of GPRC5A in diseases. Oncoscience 1, 765–776 (2014).

49. Qian, X., Jiang, C., Shen, S. & Zou, X. GPRC5A: An emerging prognostic biomarker for predicting malignancy of Pancreatic Cancer based on bioinformatics analysis. J. Cancer 12, 2010–2022 (2021).

50. Yang, L., Zhao, S., Zhu, T. & Zhang, J. GPRC5A Is a Negative Regulator of the Pro-Survival PI3K/Akt Signaling Pathway in Triple-Negative Breast Cancer. Front. Oncol. 10, 624493 (2020).

51. Chen, J., et al. Androgen receptor-regulated circ FNTA activates KRAS signaling to promote bladder cancer invasion. EMBO reports 21, e48467 (2020).

52. Tian, J., et al. circ-FNTA accelerates proliferation and invasion of bladder cancer. Oncol. Lett. 19, 1017–1023 (2020).

53. Cox, A.D. & Der, C.J. Farnesyltransferase inhibitors and cancer treatment: targeting simply Ras? Biochimica et Biophysica Acta (BBA)-Reviews on Cancer 1333, F51–F71 (1997).

54. Head, J. & Johnston, S.R. New targets for therapy in breast cancer: farnesyltransferase inhibitors. Breast Cancer Res. 6, 1–7 (2004).

55. Rowinsky, E.K., Windle, J.J. & Von Hoff, D.D. Ras protein farnesyltransferase: a strategic target for anticancer therapeutic development. J. Clin. Oncol. 17, 3631–3652 (1999).

56. Sebti, S.M. & Hamilton, A.D. Farnesyltransferase and geranylgeranyltransferase I inhibitors and cancer therapy: lessons from mechanism and bench-to-bedside translational studies. Oncogene 19, 6584–6593 (2000).

57. Fish, L., et al. Cancer cells exploit an orphan RNA to drive metastatic progression. Nat. Med. 24, 1743–1751 (2018).

58. Santhekadur, P.K. & Kumar, D.P. RISC assembly and post-transcriptional gene regulation in Hepatocellular Carcinoma. Genes Dis 7, 199–204 (2020).

59. Pan, X., Wang, Y., Lübke, T., Hinek, A. & Pshezhetsky, A.V. Mice, double deficient in lysosomal serine carboxypeptidases Scpep1 and Cathepsin A develop the hyperproliferative vesicular corneal dystrophy and hypertrophic skin thickenings. PLoS One 12, e0172854 (2017).

60. Gu, Y.Y., et al. HDAC10 Inhibits Cervical Cancer Progression through Downregulating the HDAC10-microRNA-223-EPB41L3 Axis. J. Oncol. 2022, 8092751 (2022).

61. Jiang, W. & Newsham, I.F. The tumor suppressor DAL-1/4.1B and protein methylation cooperate in inducing apoptosis in MCF-7 breast cancer cells. Mol. Cancer 5, 4 (2006).

62. Tuerxun, G., et al. Over-expression of EPB41L3 promotes apoptosis of human cervical carcinoma cells through PI3K/AKT signaling. Acta Biochim. Pol. 69, 283–289 (2022).

63. Zeng, R., et al. EPB41L3 is a potential tumor suppressor gene and prognostic indicator in esophageal squamous cell carcinoma. Int. J. Oncol. 52, 1443–1454 (2018).

64. Aoto, K., et al. ATP6V0A1 encoding the a1-subunit of the V0 domain of vacuolar H+-ATPases is essential for brain development in humans and mice. Nature communications 12, 2107 (2021).

65. Cotter, K., Stransky, L., McGuire, C. & Forgac, M. Recent Insights into the Structure, Regulation, and Function of the V-ATPases. Trends Biochem Sci 40, 611–622 (2015).

66. Capecci, J. & Forgac, M. The function of vacuolar ATPase (V-ATPase) a subunit isoforms in invasiveness of MCF10a and MCF10CA1a human breast cancer cells. J. Biol. Chem. 288, 32731–32741 (2013).

67. Wang, N., Zhu, L., Xu, X., Yu, C. & Huang, X. Integrated analysis and validation reveal ACAP1 as a novel prognostic biomarker associated with tumor immunity in lung adenocarcinoma. Comput Struct Biotechnol J 20, 4390–4401 (2022).

68. Yi, Q., Pu, Y., Chao, F., Bian, P. & Lv, L. ACAP1 Deficiency Predicts Inferior Immunotherapy Response in Solid Tumors. in Cancers (Basel), Vol. 14 (2022).

69. Zhang, J., Zhang, Q., Zhang, J. & Wang, Q. Expression of ACAP1 Is Associated with Tumor Immune Infiltration and Clinical Outcome of Ovarian Cancer. DNA Cell Biol 39, 1545–1557 (2020).

70. Lee, W.H., et al. Molecular cloning and expression of human keratinocyte proline-rich protein (hKPRP), an epidermal marker isolated from calcium-induced differentiating keratinocytes. J. Invest. Dermatol. 125, 995–1000 (2005).

71. Liu, Q., et al. Genome-wide association analysis reveals regulation of at-risk loci by DNA methylation in prostate cancer. Asian J Androl 23, 472–478 (2021).

72. Boland, M., Chourasia, A. & Macleod, K. Mitochondrial Dysfunction in Cancer. Front. Oncol. 3(2013).

73. National Cancer Institute. New Clarity on the Warburg Effect. (2022).

74. Wu, Z., et al. OMA1 reprograms metabolism under hypoxia to promote colorectal cancer development. EMBO Rep 22, e50827 (2021).

75. Hoxhaj, G. & Manning, B.D. The PI3K-AKT network at the interface of oncogenic signalling and cancer metabolism. Nat. Rev. Cancer 20, 74–88 (2020).

76. Fujii, T., et al. Implications of Low Serum Albumin as a Prognostic Factor of Long-term Outcomes in Patients With Breast Cancer. In Vivo 34, 2033–2036 (2020).

77. Soeters, P.B., Wolfe, R.R. & Shenkin, A. Hypoalbuminemia: Pathogenesis and Clinical Significance. JPEN J. Parenter. Enteral Nutr. 43, 181–193 (2019).

78. Galata, C., et al. Role of Albumin as a Nutritional and Prognostic Marker in Elective Intestinal Surgery. Can. J. Gastroenterol. Hepatol. 2020, 7028216 (2020).

79. von Meyenfeldt, M. Cancer-associated malnutrition: an introduction. Eur. J. Oncol. Nurs. 9 Suppl 2, S35–38 (2005).

80. Roux, P.P. & Topisirovic, I. Signaling Pathways Involved in the Regulation of mRNA Translation. Mol Cell Biol 38(2018).

81. Hanahan, D. Hallmarks of Cancer: New Dimensions. Cancer Discov. 12, 31–46 (2022).

82. Kern, S.E. Why your new cancer biomarker may never work: recurrent patterns and remarkable diversity in biomarker failures. Cancer Res. 72, 6097–6101 (2012).

83. Vizcaino, J.A., et al. ProteomeXchange provides globally coordinated proteomics data submission and dissemination. Nat. Biotechnol. 32, 223–226 (2014).

84. MacLean, B., et al. Skyline: an open source document editor for creating and analyzing targeted proteomics experiments. Bioinformatics 26, 966–968 (2010).

85. Krämer, A., Green, J., Pollard, J., Jr. & Tugendreich, S. Causal analysis approaches in Ingenuity Pathway Analysis. Bioinformatics 30, 523–530 (2014).

86. Liberzon, A., et al. The Molecular Signatures Database (MSigDB) hallmark gene set collection. Cell Syst 1, 417–425 (2015).

87. Yu, G., Wang, L.-G., Han, Y. & He, Q.-Y. clusterProfiler: an R Package for Comparing Biological Themes Among Gene Clusters. OMICS: A Journal of Integrative Biology 16, 284–287 (2012).

